# Conformational epistasis impairs AlphaFold structural predictions

**DOI:** 10.1101/2022.11.15.516638

**Authors:** Luciana Rodriguez Sawicki, Guillermo Benitez, Matias Carletti, Nicolas Palopoli, Maria Silvina Fornasari, Gustavo Parisi

## Abstract

Protein structures have been massively predicted using homologous sequence information. AlphaFold2 (AF2) is a recent breakthrough to predict 3D models using machine learning approaches that reached an outstanding accuracy in recent quality evaluations. However, information derived from extant homologous sequences, as those used by AF2, might not contain enough information to accurately predict protein structure. This limitation could be related to the process known as epistasis, which describes the differential effect of a mutation on the evolutionary trajectory. Clear evidence of conformational epistasis, which has a specific impact on protein structure, was characterized in the evolutionary origin of the glucocorticoid receptor (GR) specificity during its functional divergence from the mineralocorticoid (MR) receptor. In this work we explore how AF2 can reproduce conformations derived from epistatic effects. Using structural clustering and principal component analysis to analyze the structural similarities in 16 and 13 extant GR and MR conformers, respectively, we found that AF2 models for human GR failed to reproduce extant GR conformations. Interestingly, AF2 models for human MR, for which no conformational epistasis was reported, were almost indistinguishable from extant MR. Our results showcase the importance of evolutionary trajectories to predict accurate 3D models.

## Introduction

AlphaFold 2 (AF2) (Jumper et al., 2021a) has changed the way biologists study the structure-function relationship of proteins (Callaway, 2020). This foundational concept in biology breeds a myriad of areas where proteins offer a mechanistic explanation of biological processes. The AF2 breakthrough relies on a series of improvements in deep learning techniques which resulted in an unprecedented accuracy in 3D estimation as derived from the lastest CASP evaluations (Jumper et al., 2021b). In spite of its remarkable capacity in 3D estimation and its potentially powerful applications, AF2 does not resolve the protein folding problem. This long-stated puzzle is related to the understanding of how the primary structure of a protein codifies and determines the physical interactions to obtain the 3D structure of a protein (Dill and MacCallum, 2012; Dill et al., 2008). It was recently found that AF2, among other structure prediction methods, does not reproduce folding pathways and folding intermediate states (Outeiral et al., 2022). Furthermore, the AF2 output consists of top ranked models which do not reproduce the ensemble nature of proteins essential to understand protein function (Nussinov et al., 2022; Saldaño et al., 2022). The incapacity to reproduce the mechanistic processes of folding could also impair structure prediction methods due to the presence of epistatic interactions (Ortlund et al., 2007). It has been shown that epistasis during protein evolution is widespread and ubiquitous (Hinkley et al., 2011; Kryazhimskiy et al., 2011; Lehner, 2011; Lunzer et al., 2010; Starr et al., 2018). Originally used to describe the masking of one mutation by another (Bateson, 1909), the term epistasis was later applied and generalized to describe the dependence of the outcome of a mutation and the resulting phenotype on the genetic background (Phillips, 2008). For example, stabilizing substitutions could be required to occur before a slightly destabilizing one. Changing the order of these substitutions, called “permissive” mutations (Ortlund et al., 2007), can eventually destabilize the protein, producing changes in its dynamic and at last the loss of protein function. Conformational epistasis is a class of intramolecular epistasis (Lehner, 2011) in which the effect of the order of mutations impacts on the conformational changes (phenotypes) associated with the gain or loss of biological activity. A well studied system showing conformational epistasis involves the emergence of glucocorticoid specificity during the evolution of glucocorticoid receptors (Bridgham et al., 2006; Ortlund et al., 2007).

Using ancestral reconstruction and resurrection techniques (Joy et al., 2016; Thornton, 2004), Joseph Thornton’s group unveiled the evolutionary origin of GR specificity for cortisol (Bridgham et al., 2006; Ortlund et al., 2007). MR and GR evolved from a common ancestor (AncCR) by gene duplication ~470 million years ago during the early evolution of vertebrates (Figure 1A). AncCR is a promiscuous receptor activated by aldosterone, cortisol, and deoxycortisol ligands. As MR can be activated by aldosterone and to a lesser extent by cortisol, the specificity of GR for cortisol and its incapacity to be activated by aldosterone are derived characters in the GR evolutionary history. To explain the origin of GR specificity, Thornton’s group resurrected two other ancestral proteins, AncGR1 and AncGR2 (~440 and ~420 million years ago, Figure 1A and 1B) showing functional divergence but high structure conservation (RMSD between them < 1 Å). AncGR1 is also a receptor with promiscuous activation by cortisol and aldosterone like AncCR, but AncGR2 is functionally more similar to modern GRs, preferentially being activated by cortisol. There are 36 substitutions between AncGR1 and AncGR2, among which S106P and L111Q (named as group X of substitutions) are conserved in modern GRs and essential to confer high specificity for cortisol and low for aldosterone (Li et al., 2005). The substitution S106P induces a kink in the interhelical loop of AncGR2, which is evidenced in the structural alignment with AncGR1 by the large distance between Cα of P106 and S106 (Figure 1B). This substitution allows the repositioning of L111Q to form a new hydrogen bond with the dexamethasone C17-hydroxyl group (equivalent to the unique C17-hydroxyl of cortisol) (Figure 1B). The same kink is observed in modern GR when compared with MR (Figure 1C).

**Figure 1.**
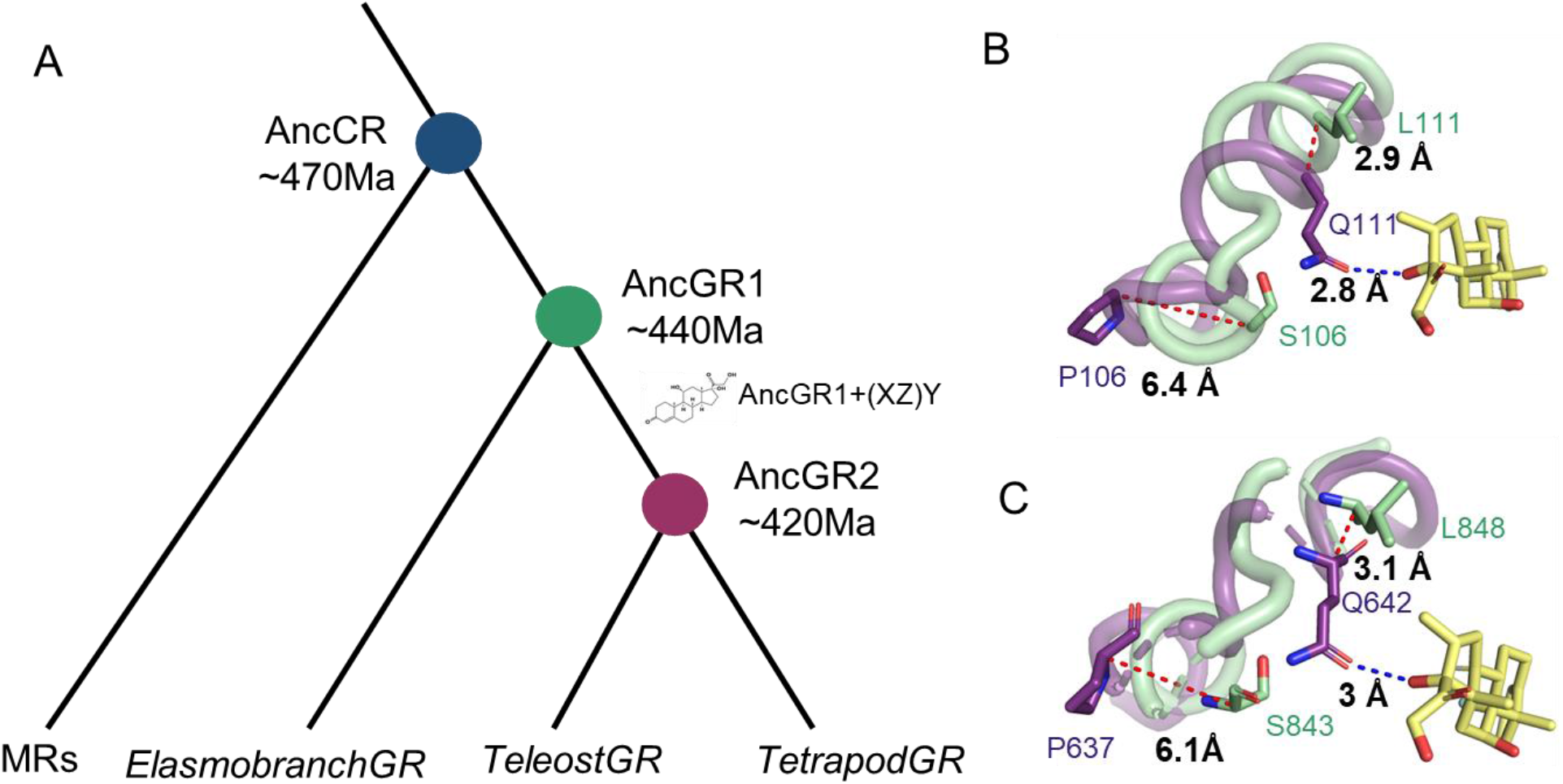
A. Functional evolution of GR and MR from a common ancestor AncCR. Evolutionary trajectory from AncGR1 with different substitutions to AncGR2, drives the evolution of cortisol specificity. AncGR2 is the common ancestor of modern GR sharing similar biological activities. B. Structural change conferring new ligand specificity. Backbone superposition of helices 6 and 7 from AncGR1 (PDB ID 3RY9, green) and from AncGR2 (PDB ID 3GN8, purple) in complex with dexamethasone (yellow). Distances between Cα atoms of S106 (AncGR1) and P106 (AncGR2), and of L111 (AncGR1) and Q111 (AncGR2), as discussed in the text, are indicated as blue dashed lines. Epistatic substitutions (groups X, Y and Z see main text) produced a new hydrogen bond between Q111 from AncGR2 and the C17-hydroxyl group of dexamethasone (red dashed line) that is absent in AncGR1. C. Backbone structure superposition of crystal structures of human MR (PDB ID 2AA2, green) and human GR (PDB ID 1M2Z, purple) bound to dexamethasone (yellow). Distances between Cα atoms of S843 (MR) and P637 (GR), and of L848 (MR) and Q642 (GR) (equivalent to P106 and Q111 in AncGR2) are indicated. The interaction between GR and dexamethasone is missing in MR. Figure adapted from Ortlund et al 2007 (Ortlund et al., 2007).

When group X substitutions were introduced to AncGR1 they produced a partial functional shift towards the biological properties of AncGR2. Thornton and co-workers discovered another two sets of substitutions that fully shift AncGR1 to the AncGR2 biological activity, which they named group Y (S212Δ, L29M and F98I) and group Z (N26T and Q105L). Moreover, they observed that one of these two sets was required to occur before the other in order to stabilize the structure, indicating a high epistatic behavior. When S106P, L111Q and the other Y and Z sets of substitutions were introduced in AncGR1 in a precise order, a full conversion of the biological properties of AncGR1 to those of AncGR2 was observed (see Figure 1A). As the conformational change associated with this shift in the biological activity highly depends on the order of the substitutions, this change is called conformational epistasis (Ortlund et al., 2007).

In this work we used human GR to test the ability of AF2 to predict protein structures which evolved under conformational epistasis. To this end we obtained AF2 models for the human GR under different hypotheses, which were later structurally compared with extant and ancestral structures of GR. We found that AF2 is unable to predict conformations that were derived from evolutionary processes under conformational epistasis. This impairment could be related with the fact that AF2 predictions are obtained with extant multiple sequence alignments which lack information about the evolutionary trajectories.

## Results

### Global structural comparisons

AF2 models for human GR (Uniprot ID P04150) and MR (Uniprot ID P08235) were obtained using the ColabFold service (Mirdita et al., 2021). GR and MR models have excellent quality as derived from the average plDDT (predicted local Distance Difference Test) scores (Mariani et al., 2013) of the 5 top models (94.73 and 95.09 in average for GR and MR models, respectively). We used the global RMSD as a dissimilarity measurement to structurally compare the five top AF2 models of GR and MR with 16 GR and 13 MR extant conformations obtained from PDB, and with the structures of AncCR, AncGR1 and AncGR2 (see Data Availability Table 1). The structural similarity clustering, based on global RMSDs, is shown in Figure 2. This clustering divides GRs and MRs into two different groups. The MRs group contains AncCR, AncGR1 and extant MRs structures, all in separate clusters. AF2 models for MR are also included in the MRs cluster, differing from most of the other structures in less than 0.5 Å, which is usually taken as the crystallographic error (Kuriyan et al., 1991). The GRs group contains three main clusters including extant GRs, AncGr2 structures and AF2 models. In this case, AF2 models differ from extant GRs with RMSD ~0.95 Å and cluster together with AncGr2.

**Figure 2.**
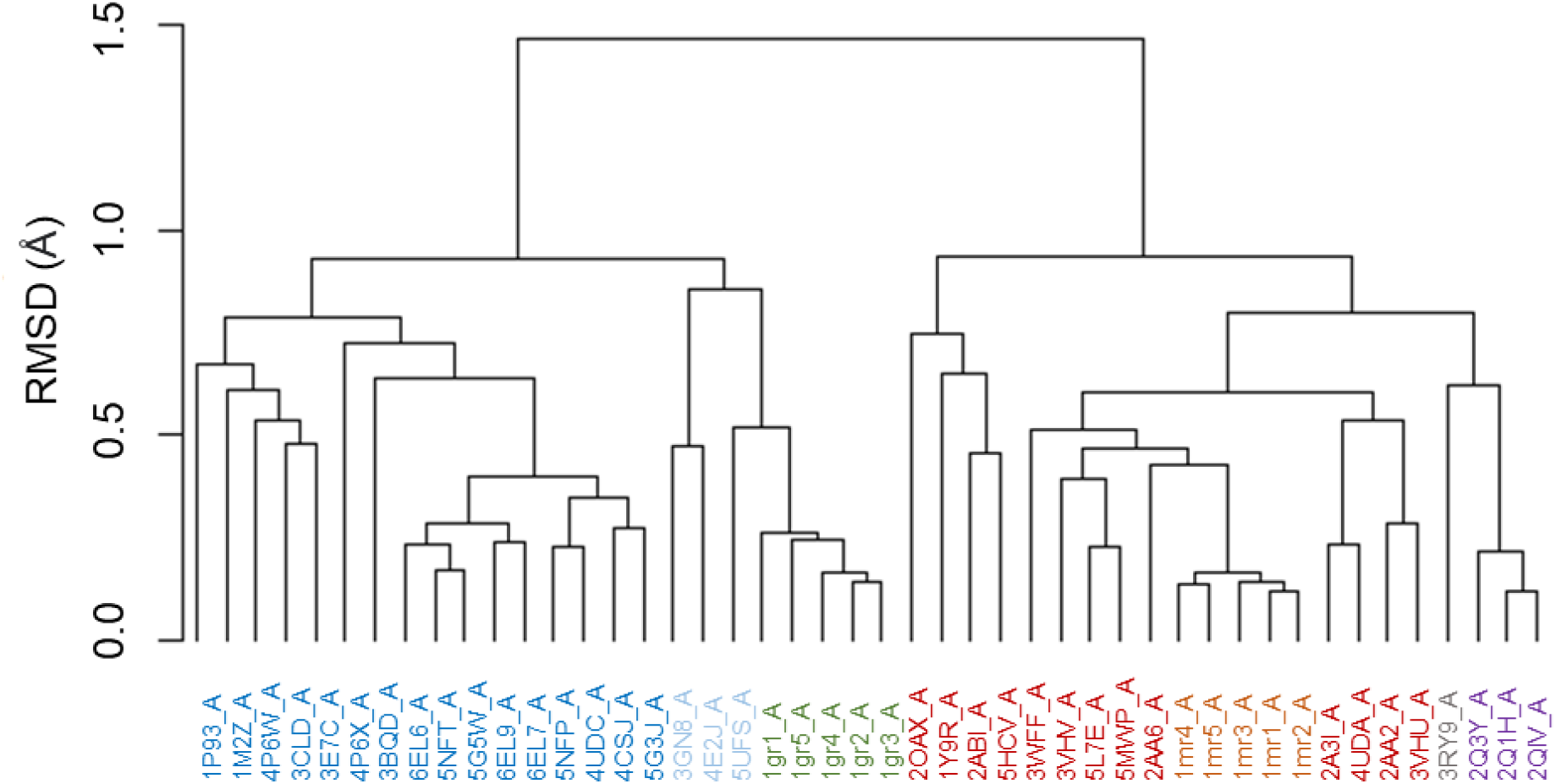
Structural clustering based on RMSD comparisons between extant GRs (blue), AncGR2s (light blue), AF2 models of GR (green), extant MRs (red), AF2 models of MR (orange), AncGR1 (grey) and AncCRs (purple).

**Table 1:**
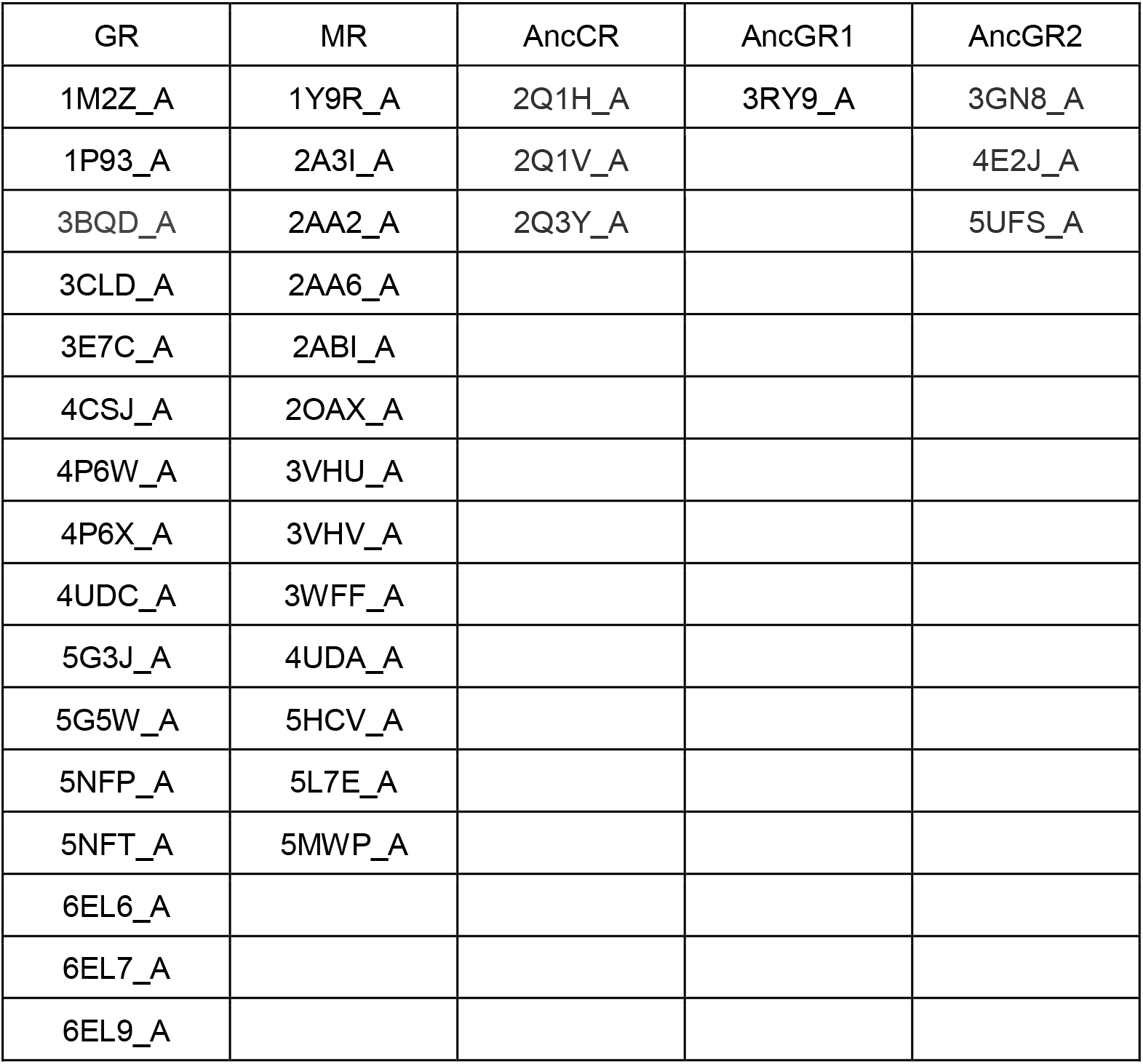
PDB IDs of GR, MR and ancestral resurrected protein structures used in this work.

The clustering perfectly fits what is expected from biological activity data. It was described that AncCR and AncGR1 share functional properties with extant MR. However, as described before, AncGR2 is more related to GRs and clusters in that group (Bridgham et al., 2006; Ortlund et al., 2007). The global RMSD comparison indicates that AF2 models for human GR are more similar to AncGR2 than to extant GRs.

To study the differences between extant GR structures and AF models we calculated the RMSD per position. We first studied the conformational diversity in human GRs to detect highly flexible regions. Figure 3A shows the RMSD per position for 16 GR conformers taken from CoDNAs database (Monzon et al., 2016) against a reference structure (PDB 1M2Z, human GR bound with dexamethasone). It is possible to see five peaks corresponding to highly flexible loops. The first region is only seen in 3BQD structure and corresponds to residues E542 to D549, which form an extended loop because of the presence of the agonist ligand (Suino-Powell et al., 2008). The GR pocket is expanded compared to the pocket observed in the dexamethasone-bound structure because of small movements in loops (regions numbered 2-5 in Figure 3A) as well as changes in side chain conformations due to binding of different ligands included with GR conformers (Biggadike et al., 2008; Kauppi et al., 2003; Ripa et al., 2018; Suino-Powell et al., 2008).

**Figure 3.**
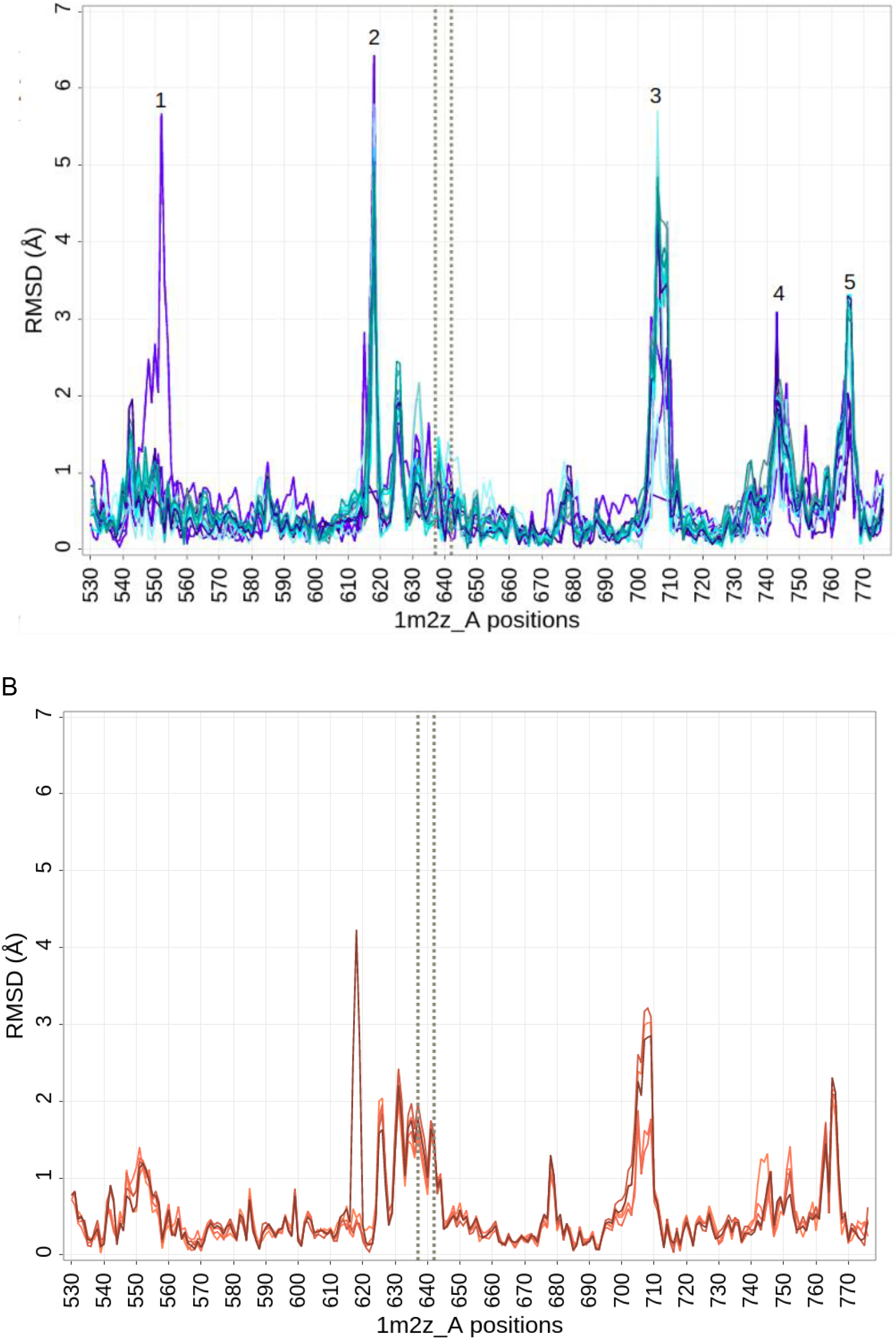
Structural comparison between structure 1M2Z_A vs the dataset of conformers of human GR (see Table 1) (A) or vs AF2 models for human GR (B). The X axis indicates 1M2Z_A positions and the Y axis shows RMSD per position in Å. Dotted lines indicate P637 and Q642 from 1M2Z.

Figure 3B shows RMSD per position between the GR reference structure and AF2 models of human GR. The AF2 models predict the same peaks that correspond to the flexible regions described above. However, AF2 models show high RMSD values around two key positions P637 and Q642 (1M2Z numeration and shown with dotted lines in Figures 3). These substitutions are near the binding site and their replacements in AncGR1 from AncGR2 (S106P, L111Q) increase cortisol specificity and highly decrease specificity for aldosterone. Basically, these historical substitutions introduce a rearrangement of an adjacent loop in helix 7 of the GR fold (Figure 1B). As we mentioned before, these substitutions have an important epistatic behavior because they should be preceded by “permissive” substitutions during evolution. As shown in Figure 2 and Figure 3B, AF2 models resemble ancestral states of GRs, with AncGR2 possibly indicating that AF is unable to reproduce epistatic interactions to properly fold extant GRs. Interestingly, AF2 models obtained using templates gave the same results as those predicted without the use of templates, both clustering with the rest of the models of GR (data not shown).

As global and position-specific Cα RMSDs only evidence backbone differences, we also measured distances between Q642 with dexamethasone co-crystallized in the reference PDB and in two other structures co-crystallized with the same ligand (PDB IDs 1M2Z, 1P93 and 4UDC, respectively), as well as in AF2 models (equivalent Q113). We found that the average distance between the carbonyl of the amide group of Q642 and the dexamethasone C17-hydroxyl group is 3.13Å in PDB structures, while in AF2 models the distance jumps to 6.24Å (see Methods). As the arrangement of Q113 (equivalent to Q642 in 1M2Z) in AF2 models could result from an artifact of AF2 modeling, we used SCWRL4 (Krivov et al., 2009) to improve the side-chain conformation of that residue. However, even after optimization of the rotamer, the distance from Q113 to dexamethasone is still higher (4.5Å) than in extant PDBs containing dexamethasone.

### Principal Component Analysis of GRs models

To further analyze structural similarities between AF2 models and extant GRs, we used principal component analysis. To improve the discrimination power, we used only the region comprising helix 7 and the loop preceding it (positions 629 to 660 in 1M2Z) described before (Figure 1C). This region also showed a high quality as derived from plDDT with an average for the top five models of 94.94. Figure 4 contains the clustering using the first two principal components (which accounts ~88% of the total structural variance) which essentially shows mostly the same clustering obtained using global RMSD in Figure 3. Again AF2 models for GRs form a different cluster from extant GRs. This means that structural differences are mostly concentrated in the region evidencing epistatic conformational variation. The only difference in reference with the RMSD clustering shown in Figure 2, is that the structure of GR 4P6W co-crystallized with mometasone furoate (MF) displaces the conformation of GR clustering in this PCA together with AF2 models and AncGR2. In the MF-bound structure (4P6W) the lipophilic C-17α furoate group fits nicely into the hydrophobic cavity and makes extensive hydrophobic interactions with the surrounding F623, I629, M639, and C643 amino acids (He et al., 2014), compared to the hydrogen bond observed between Q642 of GR and C-17 hydroxyl group from dexamethasone ligand.

**Figure 4.**
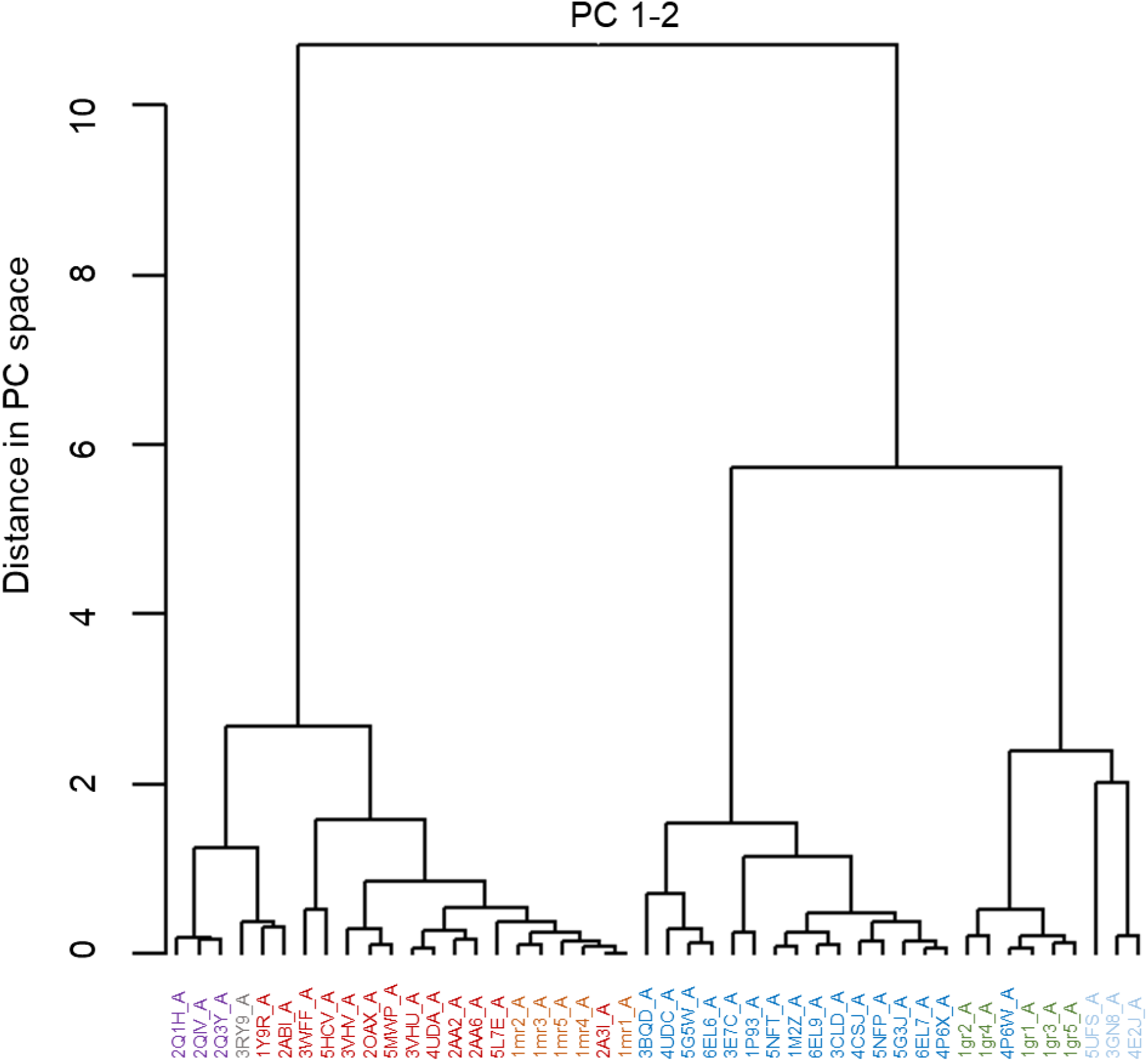
PCA clustering between extant GR (blue), AncGR2 (light blue), AF2 models for GR (green), extant MR(red), AF2 models for MR (orange), AncCR (purple) and AncGR1 (grey).

### Subsampling alignments and ancestral reconstruction

Recently different works showed results of increased capacity of AF2 to reproduce protein conformational diversity using alignment subsampling (Del Alamo et al., 2022; Wayment-Steele et al., 2022). In an attempt to increase the information of evolutionary trajectories in the input used by AF2, we used ancestral reconstruction prediction (Joy et al., 2016). Using the sequence alignment derived from AF2 homology searches in the prediction of GR structure, we estimated all the corresponding ancestral sequences using PAML (Yang, 2007) and Lazarus package (Hanson-Smith et al. 2010). We obtained AF2 models for GR using all the ancestral sequences, and using extant plus ancestral sequences in order to obtain better models. The use of ancestral sequences did not improve the quality of the models (Figure 5). All the obtained models using ancestral sequences in the alignment clustered together with the models without the ancestral reconstruction information. We also selected clusters containing the GR sequence “signature”, sequences containing S106P, L111Q and the group of Y and Z substitutions mentioned above, to derive AF2 models. In this case the GR models obtained were really poor, differing from the GR structures in RMSD’s values of ~10 A.

**Figure 5:**
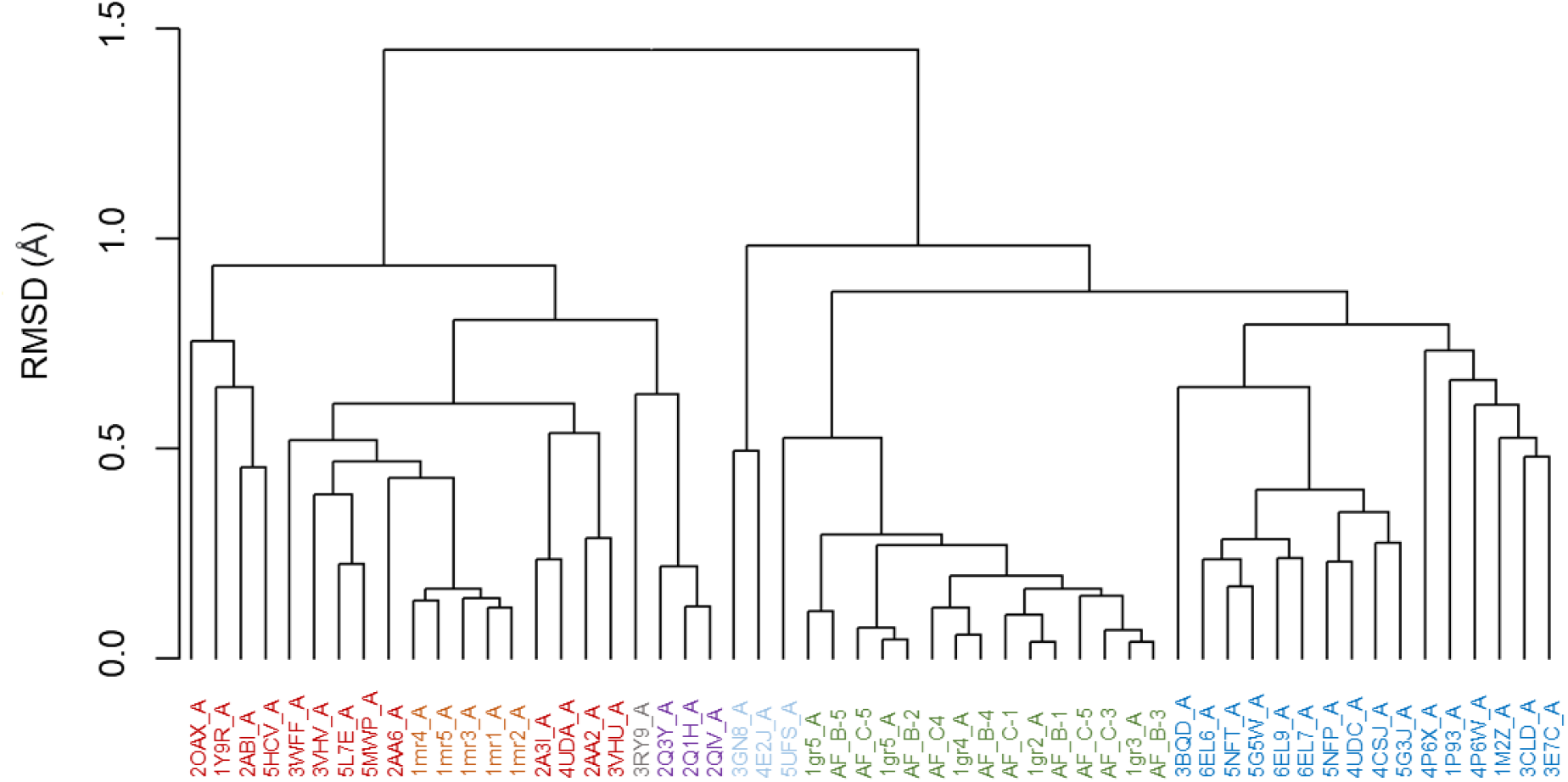
Structural clustering based on RMSD comparisons between extant GR (blue), AncGR2 (light blue), AF2 models for GR (green), extant MR(red), AF2 models for MR (orange), AncCR (purple) and AncGR1 (gray), AF2 models for GR using ancestral reconstruction sequences (AF_B 1 to 5, green) and ancestral reconstruction sequences plus extant homologous sequences (AF_C 1 to 5, green). Again, as in Figure 2, AF2 models form a different cluster from extant GR.

## Discussion

We have found that AF2 models of human GR structurally differ from extant GRs. These structural differences are concentrated in a region essential to confer the high specificity for cortisol and low for aldosterone which are the main biological characteristic of GRs. As described, this region is affected by a large epistatic effect and then is highly dependent on the trajectory or the ordered succession of substitutions during the evolutionary process. As extant structures and sequences are used by AF2 in the learning process to derive 3D models, it would be difficult to properly model epistatic interactions. This “horizontal” approach (Harms and Thornton, 2010), the use of modern sequences, has been extensively used to infer key functional related residues (Barnsley and Ondrechen, 2022; Chen et al., 2019; Donald and Shakhnovich, 2009; Melamed et al., 2015). Swapping these “key” residues between homologous proteins was not enough to fully interconvert their biological properties due to the epistatic interactions of those “key” residues (Gerlt and Babbitt, 2009; Xiang et al., 1999). In the same way, comparison of modern sequences was extensively used to predict protein structure using homology modeling (Monzon et al., 2017; Schwede et al., 2003; Wallner and Elofsson, 2005). However, the “transfer” of sequence information to a structure template should suffer the same consequences due to epistatic interactions. In fact, in very few cases, epistatic information was successfully used to improve prediction of protein structures (Rollins et al., 2019).

Our results suggest that in spite of its incredible capacity, AF2 is still a “horizontal” approach to predict protein structures. The impact of these inaccuracies in structure prediction will certainly depend on the extension of intramolecular protein sequence epistasis, which according to recent results could be ubiquitous (Lehner, 2011; Lunzer et al., 2010; Poelwijk et al., 2019).

## Materials and Methods

### AF2 models prediction

We used Colabfold (Mirdita et al., 2022) to obtain the top five AlphaFold2 3D models for Uniprot IDs P04150 (human GR) and P08235 (human MR), with and without the use of templates and always using Amber optimization.

### Structural analysis

All the extant and ancestral structures were obtained from PDB (see Data Availability Table 1). Structural alignment, pairwise RMSD calculations and RMSD-based clustering, and structural comparison by PCA analysis were performed using the R package Bio3D (Grant et al., 2006). RMSD per position was derived using Pymol. Dexamethasone was incorporated into AF2 models by structural alignment with 1M2Z_A. Distances between Q113 of AF2 models (Q642 according to PDB ID 1M2Z) and C17-hydroxyl group of dexamethasone were derived using Pymol. Different conformations for residue Q113 of AF2 models (Q642 according to PDB ID 1M2Z) were obtained using SCWRL4 (Krivov et al., 2009) for each of the top five AF2 models of human GR.

### Ancestral sequence reconstruction

Using the AF2 alignment derived for 3D prediction, consisting of 1758 homologs of human GR, we estimated a maximum likelihood phylogenetic tree with the IQ-TREE software (Nguyen et al., 2015). The tree and the mentioned alignment were used to infer the ancestral protein sequences by maximum likelihood using the PAML program (Yang, 2007). For this reconstruction, the JTT model was assumed with a discrete gamma distribution using four substitution rates among sites. Also, we used the Lazarus package for the treatment of gaps (Hanson-Smith et al., 2010). We then built two different sets of sequences as input for AF2: one containing all the ancestral sequences obtained as described above, and another set containing the ancestral sequences plus extant sequences. Both sets were aligned using ClustalO (Sievers et al., 2011) and used to obtain structural models with ColabFold.

## Acknowledgements

LRS, MSF, NP and GP are researchers and GB and MC are PhD fellows from Consejo Nacional de Investigaciones Científicas y Técnicas (CONICET). This work was supported by Universidad Nacional de Quilmes (PUNQ 1004/11 and PP 2019), Agencia Nacional de Promoción Científica y Tecnológica (PICT2018-3457, PICT 2020-0269) and the European Union’s Horizon 2020 Research and Innovation Programme (Grant Agreement N° 778247 and N° 823886). The funders had no role in study design, data collection and analysis, decision to publish, or preparation of the manuscript.

